# Analysis of flavonol regulator evolution in the Brassicaceae reveals *MYB12, MYB111* and *MYB21* duplications associated with *MYB11* and *MYB24* gene loss

**DOI:** 10.1101/2022.04.06.487363

**Authors:** Hanna M. Schilbert, Beverley J. Glover

## Abstract

**Background:** Flavonols are the largest subgroup of flavonoids, possessing multiple functions in plants including protection against ultraviolet radiation, antimicrobial activities, and flower pigmentation together with anthocyanins. They are of agronomical and economical importance because the major off-taste component in rapeseed protein isolates is a flavonol derivative, which limits rapeseed protein use for human consumption. Flavonol production in *Arabidopsis thaliana* is mainly regulated by the subgroup 7 (SG7) R2R3-MYB transcription factors MYB11, MYB12, and MYB111. Recently, the SG19 MYBs MYB21, MYB24, and MYB57 were shown to regulate flavonol accumulation in pollen and stamens. The members of each subgroup are closely related, showing gene redundancy and tissue-specific expression in *A. thaliana*. However, the evolution of these flavonol regulators inside the Brassicaceae, especially inside the Brassiceae, which include the rapeseed crop species, is not fully understood.

**Results:** We studied the SG7 and SG19 MYBs in 44 species, including 31 species of the Brassicaceae, by phylogenetic analyses followed by synteny and gene expression analyses. Thereby we identified a deep *MYB12* and *MYB111* duplication inside the Brassicaceae, which likely occurred before the divergence of Brassiceae and Thelypodieae. These duplications of SG7 members were followed by the loss of *MYB11* after the divergence of *Eruca vesicaria* from the remaining Brassiceae species. Similarly, *MYB21* experienced duplication before the emergence of the Brassiceae family, where the gene loss of *MYB24* is also proposed to have happened. Due to the overlapping spatio-temporal expression patterns of the SG7 and SG19 MYB members in *B. napus*, the loss of *MYB11* and *MYB24* is likely to be compensated by the remaining homologs.

**Conclusions:** We identified a duplication of *MYB12, MYB111*, and *MYB21* inside the Brassicaceae which is associated with *MYB11* and *MYB24* gene loss inside the tribe Brassiceae. We propose that gene redundancy and meso-polyploidization events have shaped the evolution of the flavonol regulators in the Brassicaceae, especially in the Brassiceae.

## Background

The mustard family (Brassicaceae) consists of 351 genera and almost 4000 species [1]. It contains the model plant *Arabidopsis thaliana* and several important crop plants including oilseed rape (*Brassica napus*) and cabbage (*Brassica oleracea*) domesticated for industrial use including food and biofuel production. Recent advances in Brassicaceae taxonomy revealed 51 monophyletic groups (tribes) [2, 3, 1, 4], which can be assigned to major evolutionary lineages. Around 32 million years ago (MYA) the tribe Aethionemeae diverged from the rest of the family [5]. The diversification of the other 50 tribes began ∼23 MYA and they are grouped into three [6, 7], four [8], or five lineages/clades [9, 10] (Figure 1). Three major whole-genome duplication (WGDs) events, namely At-α, At-β and At-ɣ, have occurred in the evolution of *A. thaliana* and the core Brassicaceae, which are thought to increase the genetic diversity and species radiation [11–13]. Besides these, several meso-polyploidization events have been identified inside the Brassicaceae, e.g. in the tribe Brassiceae (Figure 1) [14–16]. The whole-genome triplication (Br-α) in *Brassica* was shown to have occurred after At-α and before the radiation of the tribe Brassiceae [14–16]. Generally, polyploidization is followed by diploidization which is frequently accompanied by genome size reduction and reorganization and therefore genetic and transcriptional changes occur [17]. These changes are the basis for the “Gene Balance Hypothesis” stating that dosage-sensitive genes like transcription factors are over-retained while genes duplicated are preferentially lost after WGD events [18, 19]. It is assumed that polyploids have an adaptive advantage conferred by the availability of duplicated genes for sub- and neofunctionalization [20].

**Figure 1:**
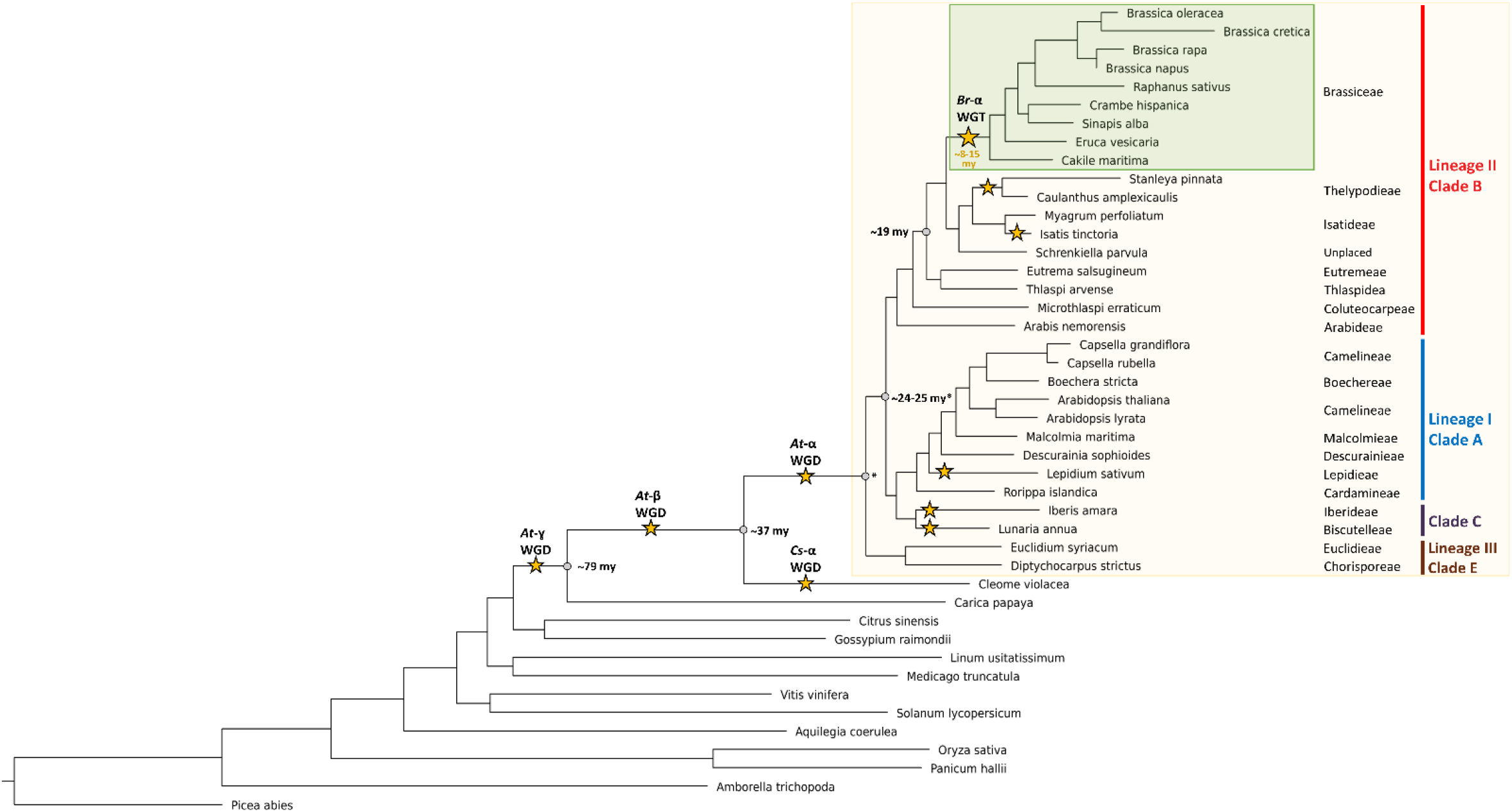
Simplified Brassicaceae phylogeny. The phylogeny of Brassicaceae family members and outgroup species is shown. The species tree was built with OrthoFinder based on proteome data sets. The Brassicaceae family is highlighted in the beige box, while species assigned to the tribe Brassiceae are highlighted in the green box. The Brassicaceae lineages and clades [9] are coloured as followed: lineage I/clade A in blue, lineage II/clade B in red, lineage III/clade E in brown and clade C in violet. Clade D is not shown as no species was analysed from this clade. Whole genome duplication (WGD) events [21–23, 4] and the Brassiceae whole genome triplication (WDT) event are marked with a star and named according to Barker *et al*., 2009. Estimated divergence times were added according to Franzke *et al*., 2011 and Walden *et al*., 2020.

One of the largest transcription factor families in plants are MYB (myeloblastosis) transcription factors [24, 25]. They play pivotal roles in regulatory networks controlling development, metabolism and responses to biotic and abiotic stresses. MYBs are classified, based on the number of up to four imperfect amino acid sequence repeats (R) in their MYB domain, into 1R-, R2R3-, 3R-, and 4R-MYBs (summarised in Dubos *et al*., 2010). Each repeat forms three a–helices. While the second and third helices build a helix–turn–helix (HTH) structure [26], the third helix makes direct contact with the major groove of the DNA [27]. There are two major models describing R2R3-MYB and R1R2R3-MYB evolution: The “loss” model states that R2R3-MYB evolved from an R1R2R3 ancestral gene by the loss of the R1 repeat [28] while the “gain” model proposes that an ancestral R2R3-MYB gene gained the R1 repeat by intragenic domain duplication leading to the emergence of R1R2R3-MYBs [29]. Recent work by Du *et al*. suggests that the gain model provides a more parsimonious and reasonable explanation for the phylogenetic distribution of two and three repeat MYBs as both MYB classes are proposed to have coexisted in primitive eukaryotes [30]. However, Jiang *et al*. inferred that the gain model is unlikely, based on phylogenetic analyses [31].

R2R3-MYBs are the largest class of MYB transcription factors as they are exceptionally expanded in plant genomes [30, 31]. For example, R2R3-MYBs account for 64% and 63% of all MYB proteins in *A. thaliana* and *B. napus*, respectively [25, 24, 32] (Figure 2). The expansion of the R2R3-MYB family in plants resulted in a wide functional diversity of R2R3-MYBs, which regulate mainly plant-specific processes like stress responses, development and specialized metabolism [24]. R2R3-MYBs can be further classified into 23 subgroups by characteristic amino-acid motifs in the C-terminal region [25]. Several subgroups are involved in the regulation of flavonoid biosynthesis, one of the best studied plant biosynthesis pathways [33]. Flavonoids are responsible for plant pigmentation and can provide protection against biotic and abiotic stresses like UV-radiation [33]. While the subgroup 6 (SG6) family members MYB75/PAP1, MYB90/PAP2, MYB113, and MYB114 regulate anthocyanin accumulation [34, 35], the SG5 member MYB123/TT2 controls proanthocyanidin biosynthesis in *A. thaliana* [36].

**Figure 2:**
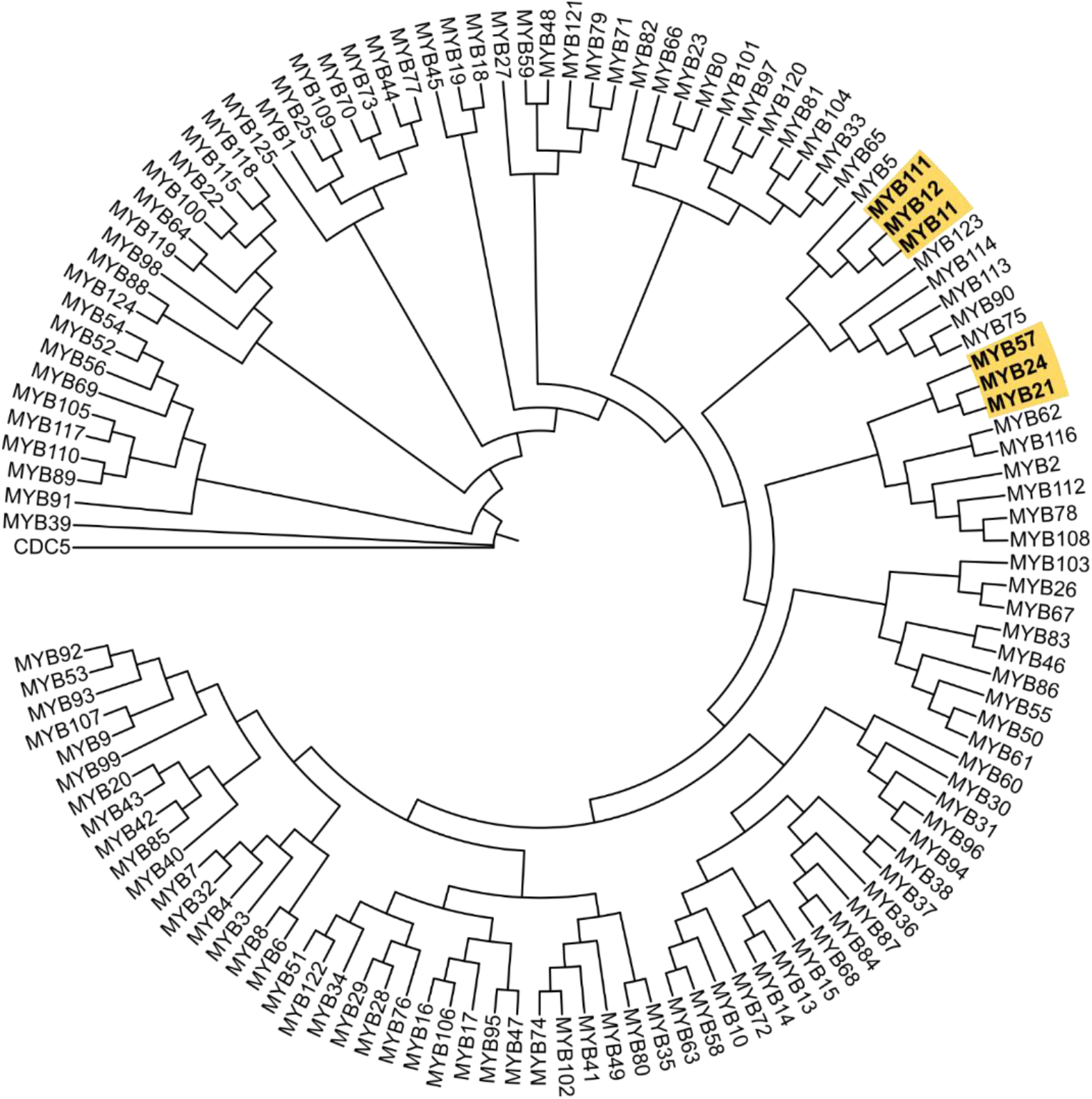
Schematic overview of the R2R3-MYB phylogeny of *Arabidopsis thaliana*. The subgroup 7 MYBs (MYB11, MYB12, MYB111) and subgroup 19 MYBs (MYB21, MYB24, MYB57) are shown in bold highlighted in yellow. The full amino acid sequences were aligned with ClustalW [53]. MEGA version 11.0.11 [54] was used to perform neighbor-joining tree analysis with 1,000 bootstraps.

Flavonols are the largest subgroup of flavonoids, and are involved in UV-protection and flower pigmentation together with anthocyanins [37, 38]. Moreover they are of agronomical and economical importance as the major off-taste component in rapeseed protein isolates is a flavonol derivative - this limits rapeseed protein palatability and human consumption [39]. The main regulators of flavonol biosynthesis in *A. thaliana* are the SG7 members MYB12, MYB11, and MYB111 [40, 41]. The SG7 MYBs show spatio-differential gene expression patterns in *A. thaliana* seedlings: *MYB12* is expressed in roots, while *MYB111* is expressed in cotyledons and *MYB11* is marginally expressed in specific domains of the seedling including the apical meristem, the primary leaves, the apex of cotyledons, at the hypocotyl–root transition, the origin of lateral roots and the root tip as well as the vascular tissue of lateral roots [41]. However, the *A. thaliana myb11/myb12/myb111* triple mutant retained flavonols in pollen grains and siliques/seeds [42]. This MYB11-, MYB12-, and MYB111-independent accumulation of flavonol glycosylates was recently addressed by the finding of a new group of flavonol regulators belonging to SG19: MYB21, MYB24, and MYB57 [43–45]. The three SG19 MYBs have previously been described to be involved in jasmonate-dependent regulation of stamen development and are expressed in all four whorls of the flower [46–48]. All SG7 MYBs can act as independent transcription factors by regulating e.g. the expression of flavonol synthase (FLS) [40, 41, 49], which produces flavonols from dihydroflavonols [50]. Studies have now shown that the SG19 MYBs can also bind and activate the *FLS1* promoter [43–45]. Moreover, MYB99 is postulated to act in a MYB triad with MYB21 and MYB24 to regulate flavonol biosynthesis in anthers [43]. The bZIP transcription factor HY5 is required for *MYB12* and *MYB111* activation under UV-B and visible light in *A. thaliana*, while MYB24 was recently shown to regulate and bind to the HYH (HY5 ortholog) promoter in *Vitis vinifera* [51, 52].

In this study we used 44 species, of which 31 belong to the Brassicaceae family, to analyse the evolution of the flavonol regulators, namely the SG7 and SG19 MYBs. In total, these 31 Brassicaceae species span 17 tribes and represent all three major lineages of the core Brassicaceae. By incorporating phylogenetic and synteny information, a duplication of *MYB12, MYB111*, and *MYB21* inside the Brassicaceae accompanied by the loss of *MYB11* and *MYB24* inside the Brassiceae was identified. Gene expression analyses suggest that gene redundancy might have played a role in the loss of *MYB11* and *MYB24*. Moreover, the meso-polyploidization events in the Brassicaceae likely shaped the evolution of flavonol regulators, especially in the tribe Brassiceae.

## Results

In this study we used a comprehensive data set collection derived from 44 species, including 31 Brassicaceae species spanning 17 tribes (Figure 1, Additional file 1). The inferred species tree revealed that most of the analysed Brassicaceae tribes are monophyletic and can be assigned to the three major lineages characteristic for the Brassicaceae family (Figure 1). In this analysis the Brassiceae tribe is represented by 9 species (*Brassica oleracea, Brassica cretica, Brassica rapa, Brassica napus, Raphanus sativus, Crambe hispanica, Sinapis alba, Eruca vesicaria, Cakile maritima*), which has the Isatideae and Thelypodieae as sister clades. The quality assessment revealed that the majority of the 44 proteome data sets (Brassicaceae and non-Brassicaceae) are suitable for this analysis due to often more than 90% complete BUSCOs (Additional file 1). The 31 Brassicaceae data sets revealed 71.2% (*Stanleya pinnata*) to 99.3% (*A. thaliana*) complete BUSCOs emphasizing the overall high completeness of these data sets.

The genome-wide identification of MYB proteins revealed different numbers of 1R-, R2R3-, 3R-MYBs and MYB-related proteins per species, ranging inside the Brassicaceae from 1 to 17 for 1R-, 90 to 442 for R2R3-, and 3 to 19 for 3R-MYBs (Additional file 2). In order to analyse the SG7 and SG19 R2R3-MYBs in the Brassicaceae in detail all respective homologs per species were extracted and used for phylogenetic analyses (Additional file 3, Figure 3). In addition, all MYB123 (SG5) and MYB99 homologs were incorporated because MYB123 regulates a competing branch of the flavonoid pathway and is sister clade to SG7, and MYB99 is proposed to act in a regulatory triad with the SG19 MYBs. Interestingly, divergence into *MYB11* and *MYB12*, as well as *MYB21* and *MYB24*, was specifically observed for Brassicaceae members, while *Cleome violacea* revealed only one *MYB11-MYB12* and *MYB21-MYB24* homolog. Additional *MYB11-MYB12* and *MYB21-MYB24* homologs from several non-Brassicaceae species like tomato were identified as clusters preceding the divergence of the Brassicaceae *MYB11, MYB12, MYB21* and *MYB24* homologs. This suggests the emergence of separate *MYB11* and *MYB12* as well as *MYB21* and *MYB24* clades after the divergence of the Cleomaceae from its sister group the Brassicaceae (Figure 3).

**Figure 3:**
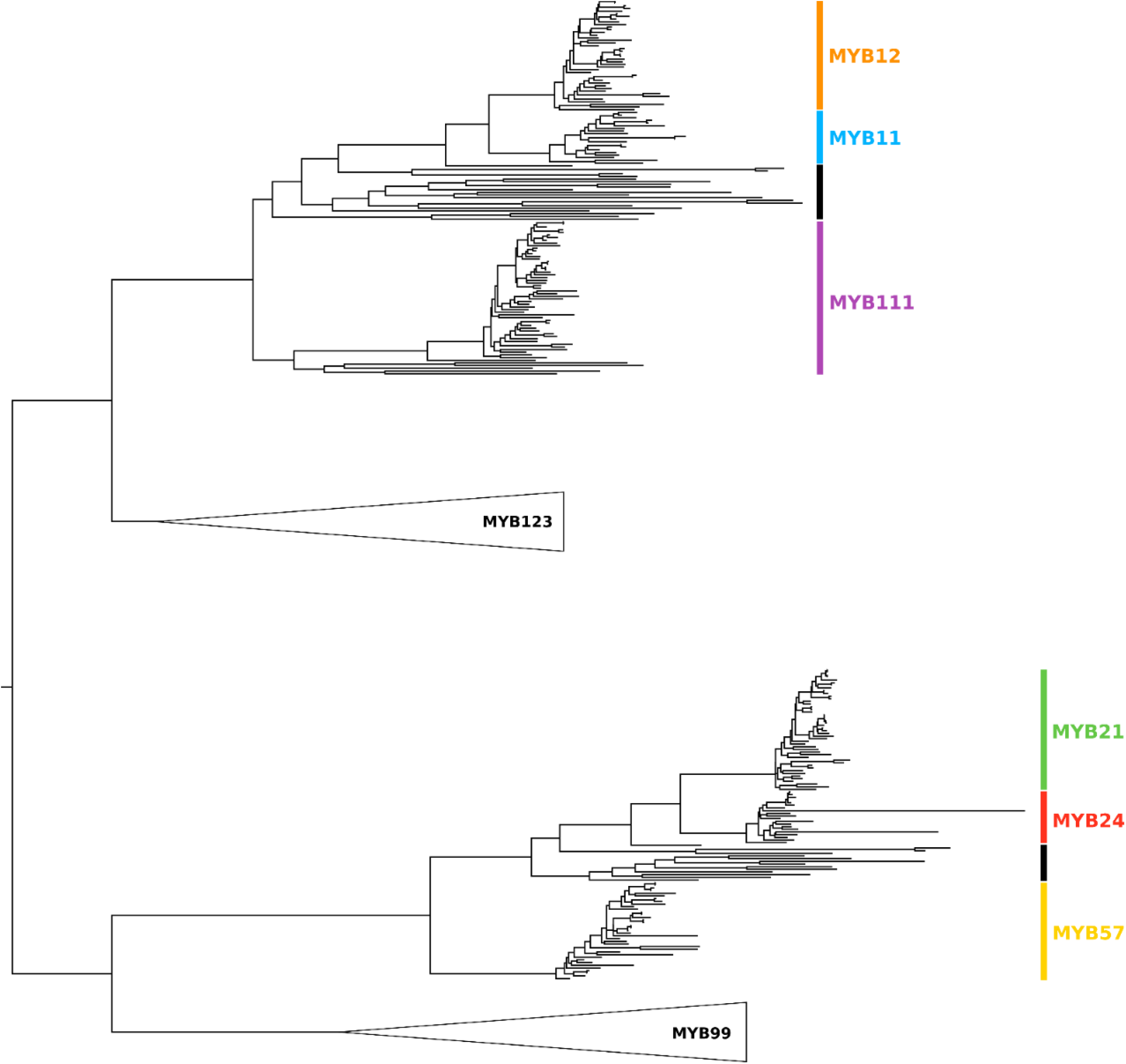
Scheme of the phylogenetic relationships of SG7 and SG19 members. The phylogenetic relationship of the SG7 (*MYB11, MYB12, MYB111*) and SG19 MYBs (*MYB21, MYB24, MYB57*) is displayed. The classification per clade is based on the respective *A. thaliana* homolog: the *MYB12* clade is coloured in orange, *MYB11* in light blue, *MYB111* in violet, *MYB21* in green, *MYB24* in red, and *MYB57* in yellow. The black vertical bars inside the SG7 and SG19 clades mark the *MYB11-MYB12* and *MYB21-MYB24* sequences derived from species outside of the Brassicaceae, respectively. The *MYB123* and *MYB99* clade were collapsed and are represented by triangles as labeled. The figure is not to scale.

### Phylogeny of SG7 MYBs

The phylogenetic analysis of SG7 members *MYB11, MYB12*, and *MYB111* revealed that at least one *MYB111* homolog is present per Brassicaceae species, except for *Arabis nemorensis* (Figure 4, Additional file 3, Additional file 4). Similarly, the majority of Brassicaceae members contained one *MYB12* homolog. However, all Brassiceae species possess a duplication of *MYB12* and *MYB111* (Figure 4). At least two *MYB111* and *MYB12* homologs were also identified in the closely related species *Caulanthus amplexicaulis* and *Isatis tinctoria*, while only two *MYB111* and no *MYB12* homolog were detected in *Stanleya pinnata*. However, the duplication event in *I. tinctoria* is likely associated with the independent meso-polyploidization event occurring in this tribe as shown by the close phylogenetic relationship of the respective *MYB111* and *MYB12* homologs (Figure 1, Figure 4). Even though independent meso-polyploidization events have also occurred in *C. amplexicaulis* and *S. pinnata*, the respective *MYB111* homologs fall into two separate clades indicating a deeper *MYB111* duplication preceding the divergence of the Brassiceae. The same applies for the *MYB12* duplication of *C. amplexicaulis*. Interestingly, no *MYB11* homolog was identified in the *Brassica* species, *R. sativus, C. hispanica*, and *S. alba*, indicating that *MYB11* might be absent in these species (Figure 4). As two *MYB11* homologs were found in *E. vesicaria* and one in *C. maritima*, this gene loss is assumed to have occurred after the divergence of *E. vesicaria*. Moreover, no *MYB11* homolog was detected in *S. pinnata, Schrenkiella parvula, Thlaspi arvense, Malcolmia maritima, Descurainia sophioides*, and *Lepidium sativum*.

**Figure 4:**
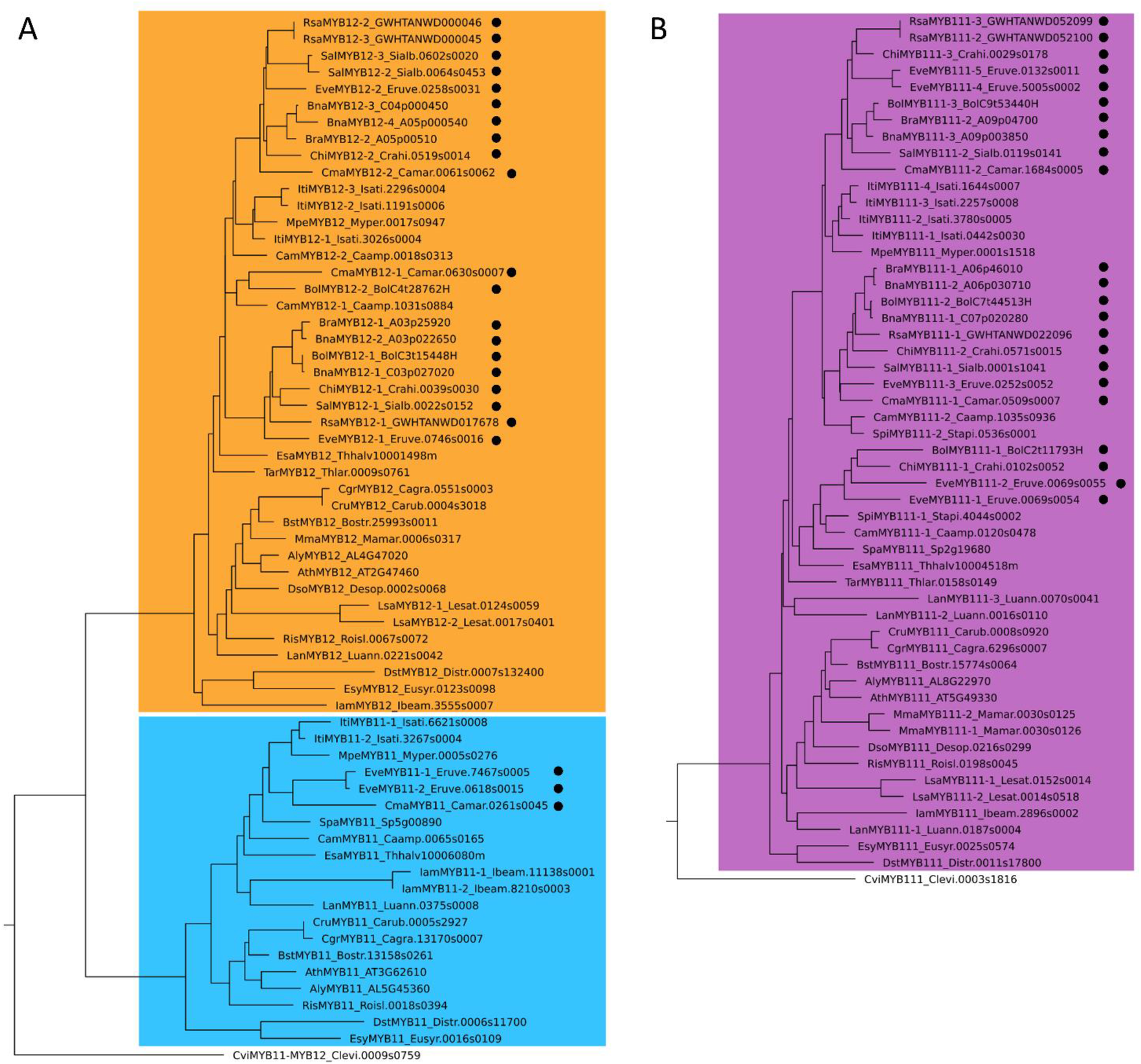
Phylogeny of SG7 members in Brassicaceae. The phylogenies of *MYB11* and *MYB12* (A) and *MYB111* (B) homologs of the Brassicaceae are displayed. Homologs of Brassiceae species are marked with a black circle. The *MYB12* clade is coloured in orange, the *MYB11* clade in light blue, and the *MYB111* clade in violet. The classification per clade is based on the respective *A. thaliana* homologs. The identified SG7 homologs of *Cleome violacea* are displayed as *C. violacea* serves as representative of the Cleomaceae, which is sister group to Brassicaceae. The figure is not to scale.

### Synteny analysis of SG7 MYBs

The potential *MYB11* gene loss inside the Brassiceae was analysed in detail by examining the degree of local synteny at the *MYB11* locus. In line with the phylogenetic analysis, *MYB11* was absent from the genomic regions of *B. napus, B. oleracea, B. rapa, R. sativus, C. hispanica*, and *S. alba* showing the highest local synteny with the corresponding *MYB11* locus from *A. thaliana*, while a *MYB11* homolog was identified for *E. vesicaria, C. maritima, I. tinctoria*, and *Myagrum perfoliatum* (Figure 5). Supporting these findings, no *MYB11* homolog was identified via a TBLASTN search against these syntenic regions, as well as the genome sequences of the *Brassica* species, *R. sativus, C. hispanica*, and *S. alba*.

**Figure 5:**
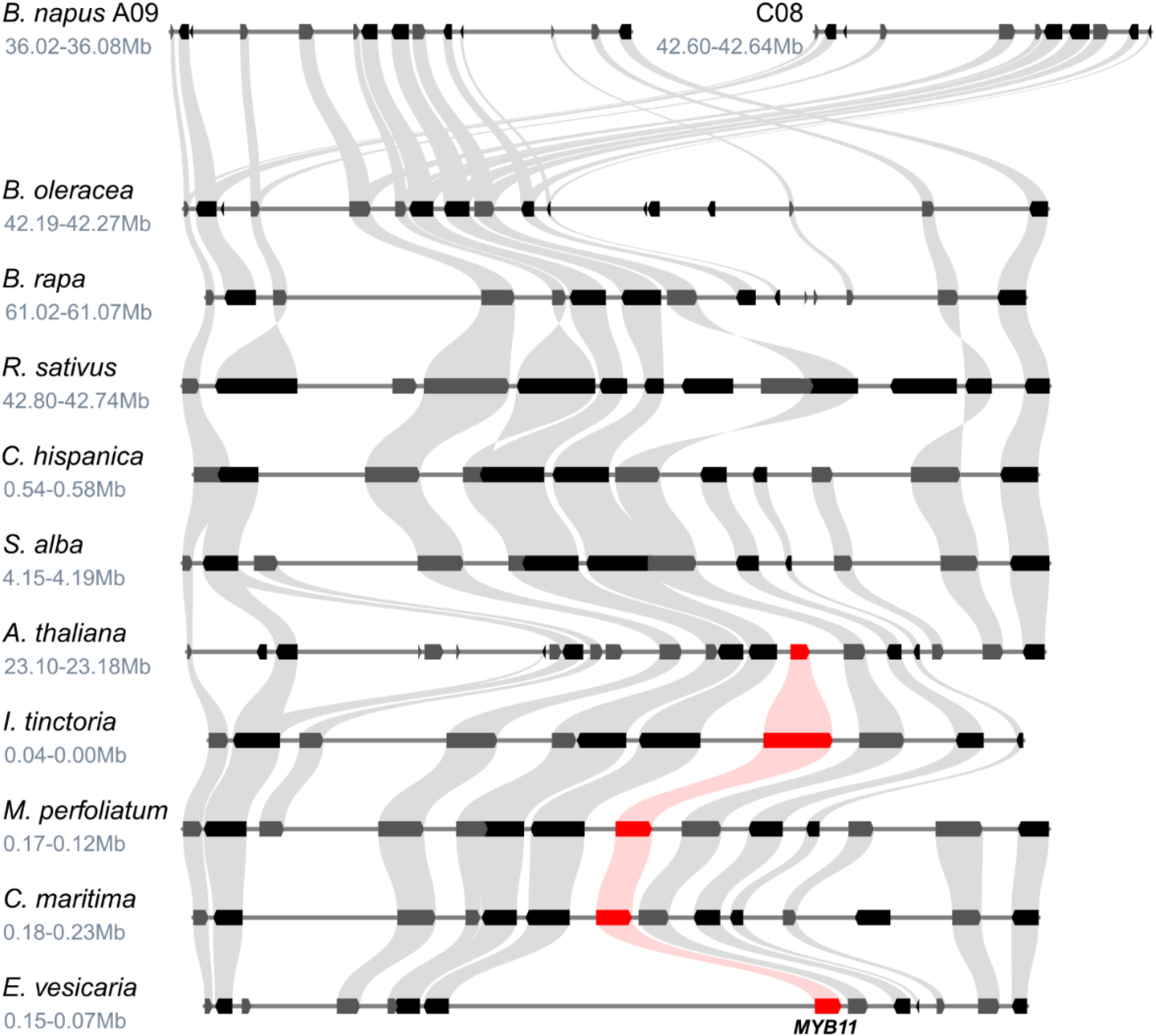
Synteny analysis of the *MYB11* locus suggests gene loss inside the Brassiceae. The syntenic relationship at the *MYB11* locus is shown for several Brassicaceae members. The position of the genomic region in the respective genome assembly is given underneath the species name in million base pairs (Mb). Grey curved beams connect the identified syntenic genes. The rectangle-shaped arrows represent annotated genes. Genes located on the forward strand are shown in grey and genes located on the reverse strand are shown in black. *MYB11* homologs are marked in red and connected by light red lines.

### Gene expression analyses of SG7 MYBs

In order to analyse the expression patterns of SG7 members in Brassiceae and to investigate whether the duplications of *MYB12* and *MYB111* result in different tissue-specific expression patterns, we harnessed RNA-Seq data sets of *B. napus* (Table 1). In general, *BnaMYB111-2_A06p030710* and *BnaMYB111-1_C07p020280* show a similar expression pattern across multiple tissues (anther, petal, bud, and silique). However, *BnaMYB111-2_A06p030710* revealed unique expression in developing seeds, seed coat, and sepals. *BnaMYB111-3_A09p003850* was not expressed in any of the analysed tissues. While all four *BnaMYB12* homologs are expressed in reproductive tissues (anthers, pistils, ovules, buds, young seeds), only three homologs (*BnaMYB12-3_C04p000450, BnaMYB12-2_A03p022650, BnaMYB12-1_C03p027020*) are additionally expressed in mature seeds and seed coat. Uniquely tissue-specific expression comparing all SG7 MYBs was identified for *BnaMYB12-3_C04p000450* in late seed coat development (35 DAF) and *BnaMYB111-2_A06p030710* is uniquely expressed in sepals and mature seeds compared to the other *BnaMYB111* homologs.

**Table 1:** Tissue-specific expression of SG7 MYBs in *B. napus*. The tissue-specific expression of the identified *MYB12* and *MYB111* homologs in *B. napus* is presented in mean transcripts per million (TPMs). The number of analysed data sets per tissue is stated in brackets (n=X). Intensity of the blue colouration indicates the expression strength (darker = stronger expression). Abbreviations: weeks after pollination (WAP), days after pollination (DAP), days after flowering (DAF), days (D), shoot apical meristem (SAM).

|  | *MYB12-1*<br>*C03p027020* | *MYB12-2*<br>*A03p022650* | *MYB12-3*<br>*C04p000450* | *MYB12-4*<br>*A05p000540* | *MYB111-1*<br>*C07p020280* | *MYB111-2*<br>*A06p030710* | *MYB111-3*<br>*A09p003850* |
| --- | --- | --- | --- | --- | --- | --- | --- |
| **SAM (n=16)** | 0.4 | 0.6 | 0.5 | 0.0 | 0.1 | 0.1 | 0.1 |
| **Anther prophase 1 (n=12)** | 2.0 | 2.9 | 1.3 | 1.6 | 19.4 | 23.5 | 0.0 |
| **Anther bolting (n=6)** | 0.3 | 0.2 | 0.6 | 0.7 | 2.9 | 2.9 | 0.0 |
| **Anther flowering (n=4)** | 1.0 | 3.0 | 3.8 | 0.6 | 5.5 | 9.1 | 0.0 |
| **Stamen (n=1)** | 0.1 | 0.1 | 0.4 | 1.0 | 0.0 | 0.0 | 0.0 |
| **Ovule (n=1)** | 4.7 | 3.2 | 3.1 | 1.5 | 0.0 | 0.8 | 0.1 |
| **Pistil (n=3)** | 0.8 | 1.5 | 1.8 | 1.1 | 0.1 | 0.3 | 0.0 |
| **Sepal (n=1)** | 0.0 | 0.0 | 0.0 | 0.0 | 0.0 | 2.3 | 0.0 |
| **Petal (n=2)** | 0.6 | 3.1 | 6.8 | 6.3 | 1.0 | 1.0 | 0.0 |
| **bud (n=33)** | 2.2 | 4.0 | 3.1 | 1.9 | 9.4 | 13.3 | 0.1 |
| **Silique 10-20DAF (n=13)** | 0.9 | 1.3 | 0.6 | 0.3 | 0.3 | 0.7 | 0.0 |
| **Silique 25DAF (n=6)** | 1.1 | 1.6 | 1.3 | 0.3 | 0.4 | 0.4 | 0.1 |
| **Silique 30DAF (n=6)** | 1.0 | 0.9 | 0.5 | 0.1 | 1.3 | 1.3 | 0.1 |
| **Silique 40DAF (n=2)** | 0.2 | 0.1 | 0.0 | 0.0 | 0.1 | 0.1 | 0.0 |
| **Seed 2WAP (n=1)** | 6.3 | 4.5 | 2.9 | 5.7 | 0.0 | 1.5 | 0.0 |
| **Seed 4WAP (n=1)** | 4.6 | 4.0 | 3.4 | 0.6 | 0.0 | 2.2 | 0.0 |
| **Seed 6WAP (n=1)** | 0.2 | 0.0 | 0.0 | 0.0 | 0.0 | 2.6 | 0.0 |
| **Seed 8WAP (n=1)** | 0.0 | 0.3 | 0.0 | 0.0 | 0.0 | 0.0 | 0.0 |
| **Seed brown 26DAF (n=1)** | 4.7 | 4.8 | 5.3 | 0.8 | 0.0 | 0.5 | 0.3 |
| **Seed yellow 26DAF (n=1)** | 3.9 | 3.5 | 4.7 | 0.3 | 0.0 | 1.6 | 0.0 |
| **Seed coat 14DAF (n=7)** | 4.8 | 6.2 | 7.2 | 0.0 | 0.0 | 1.6 | 0.0 |
| **Seed coat 21DAF (n=6)** | 7.3 | 6.7 | 17.1 | 0.0 | 0.7 | 13.8 | 0.0 |
| **Seed coat 28DAF (n=6)** | 4.2 | 4.3 | 17.1 | 0.0 | 0.1 | 0.6 | 0.0 |
| **Seed coat 35DAF (n=6)** | 0.9 | 0.4 | 6.3 | 0.0 | 0.0 | 0.2 | 0.0 |
| Seed coat 42DAF (n=6) | 0.5 | 0.1 | 0.8 | 0.0 | 0.0 | 0.1 | 0.0 |
| Embryo (n=6) | 0.8 | 0.6 | 0.0 | 0.0 | 1.3 | 2.6 | 0.2 |
| Endosperm (n=8) | 0.6 | 0.1 | 1.0 | 0.1 | 0.0 | 0.1 | 0.0 |
| Seedling (n=9) | 1.1 | 0.9 | 0.7 | 0.1 | 0.3 | 1.8 | 0.1 |
| Cotyledon 7-10D (n=34) | 0.2 | 0.3 | 0.1 | 0.0 | 0.0 | 0.2 | 0.0 |
| Leaf juvenile (n=12) | 0.7 | 0.9 | 0.7 | 0.3 | 0.5 | 0.9 | 0.0 |
| Leaf old (n=12) | 0.6 | 0.7 | 0.6 | 0.2 | 0.1 | 0.0 | 0.0 |
| Internode flowering (n=6) | 0.2 | 0.1 | 0.3 | 0.1 | 0.0 | 0.1 | 0.0 |
| Stem (n=19) | 1.8 | 2.4 | 0.6 | 0.1 | 0.2 | 0.3 | 0.0 |
| Shoot (n=2) | 0.7 | 0.5 | 1.2 | 0.5 | 0.6 | 0.4 | 0.1 |
| Shoot apices (n=2) | 0.2 | 0.8 | 1.6 | 0.8 | 0.5 | 0.2 | 0.0 |
| Root seedling (n=13) | 0.1 | 0.1 | 0.1 | 0.0 | 0.0 | 0.0 | 0.0 |
| Root 30DAP (n=20) | 0.1 | 0.1 | 0.0 | 0.0 | 0.0 | 0.0 | 0.0 |
| Root 60DAP (n=2) | 0.1 | 0.0 | 0.0 | 0.0 | 0.0 | 0.0 | 0.0 |

Three of the four *BnaMYB12* homologs (*BnaMYB12-3_C04p000450, BnaMYB12-2_A03p022650, BnaMYB12-4_A05p000540*) had overlapping co-expression patterns with genes related to flavonol biosynthesis, including *F3’H* and the flavonol glycosyltransferase *UGT84A2* (Additional file 5). However, only *BnaMYB12-1_C03p027020* and *BnaMYB12-3_C04p000450* were additionally co-expressed with *CHS, F3H, CHIL*, and *FLS1*. Interestingly, *BnaMYB12-4_A05p000540* was found to be co-expressed with *MYB106*, a transcription factor involved in trichome branching regulation in *A. thaliana*. No co-expressed genes were identified for the marginally expressed *BnaMYB111-3_A09p003850*. However, the other two *BnaMYB111* homologs were co-expressed with genes derived from the flavonoid/flavonol biosynthesis and phenylpropanoid pathway including *FLS1, F3H*, flavonol glycosyltransferases, and *4CL3* (Additional file 5).

### Phylogeny of SG19 MYBs

At least one *MYB57* and one *MYB21* homolog was identified in the analysed Brassicaceae species via phylogenetic analysis, except no *MYB57* homolog was detected in *S. pinnata* (Figure 6, Additional file 3, Additional file 4). All Brassiceae species, *C. amplexicaulis* and *I. tinctoria* revealed the presence of two *MYB21* homologs, indicating a duplication event. The *MYB21* duplication event in *I. tinctoria* is likely associated with the independent meso-polyploidization event occurring in this tribe as shown by the close phylogenetic relationship of the *MYB21* homologs (Figure 1, Figure 6). However, the *MYB21* homologs of *C. amplexicaulis* fall into two separate clades indicating a deeper *MYB21* duplication preceding the divergence of the Brassiceae. Additionally, most Brassiceae species contained two *MYB57* homologs with *C. hispanica* and *S. alba* being the exceptions with only one *MYB57* homolog identified in each of them. Besides *I. tinctoria* none of the closest sister tribes of the Brassiceae revealed more than one *MYB57* homolog. The independent meso-polyploidization event of *I. tinctoria* likely resulted in two *MYB57* homologs from which a third *MYB57* homolog likely emerged from tandem duplication. Thus, the *MYB57* duplication event likely took place after the divergence of the Brassiceae and *C. hispanica*, and *S. alba* subsequently lost one *MYB57* homolog.

**Figure 6:**
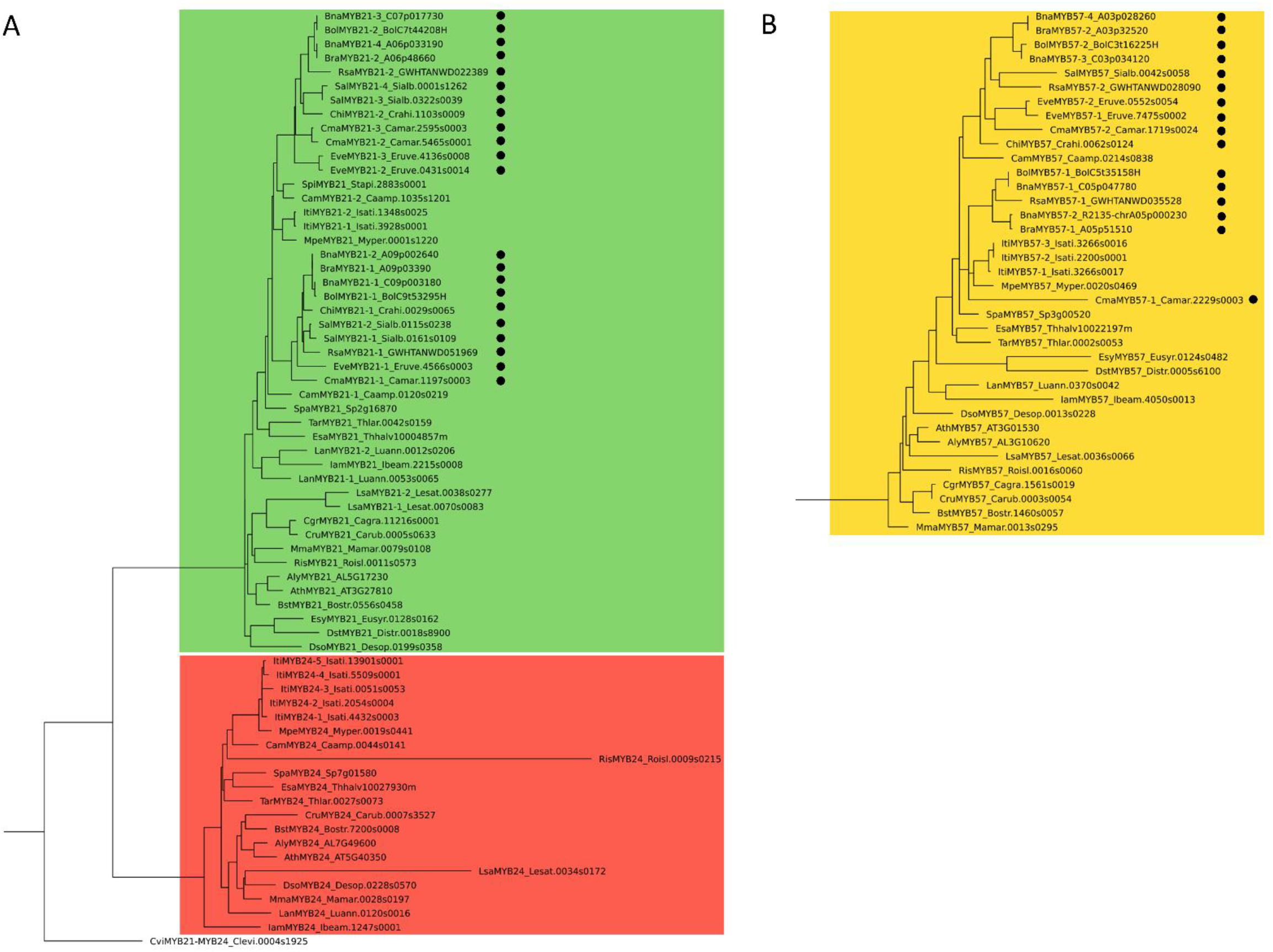
Phylogeny of SG19 members in Brassicaceae. The phylogenies of *MYB21* and *MYB24* (A) and *MYB57* (B) homologs of the Brassicaceae is displayed. The *MYB21* clade is coloured in green, the *MYB24* clade in red, and the *MYB57* clade in yellow. Homologs of Brassiceae species are marked with a black circle. The classification per clade is based on the respective *A. thaliana* homologs. The identified SG19 homologs of *Cleome violacea* are displayed as *C. violacea* serves as representative of the Cleomaceae, which is sister group to Brassicaceae. The figure is not to scale.

No *MYB24* homolog was identified in all analysed Brassiceae species, as well as *S. pinnata, A. nemorensis, Capsella grandiflora, Euclidium syriacum*, and *Diptychocarpus strictus* (Figure 6). At least one *MYB24* copy was detected in the remaining 17 Brassicaceae species. As all species of the closest Brassiceae sister tribes contain a *MYB24* homolog except for *S. pinnata*, which has a low-quality data set, the loss of *MYB24* is suggested to have occurred after the divergence of the Brassiceae tribe. Moreover, *MYB24* might have been lost in the common ancestor of *E. syriacum* and *D. strictus*.

### Synteny analysis of SG19 MYBs

In accordance with the phylogenetic analyses, *MYB24* could not be detected via local synteny analysis in *B. napus, B. oleracea, B. rapa, R. sativus*, and *S. alba*, while the locus containing a *MYB24* homolog of *M. perfoliatum* showed high local synteny to the *MYB11* locus of *A. thaliana* (Figure 7). Supporting these findings, no *MYB24* homolog was identified in the syntenic regions of *B. napus, B. oleracea, B. rapa, R. sativus*, and *S. alba* via a TBLASTN search. Additionally, no *MYB24* homolog was detected in all nine Brassiceae genome sequences.

**Figure 7:**
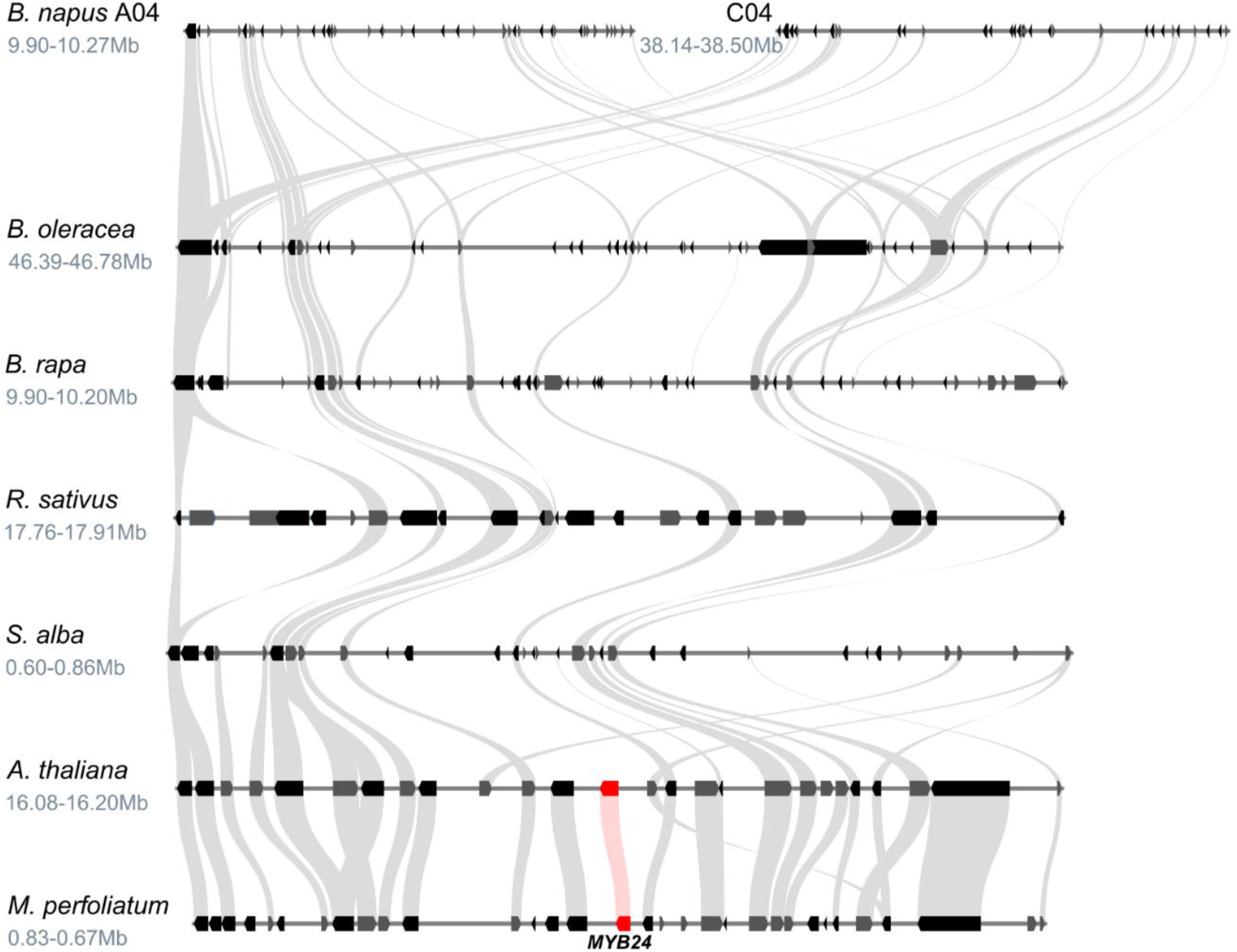
Synteny analysis of the *MYB24* locus suggests gene loss in the Brassiceae. The syntenic relationship at the *MYB24* locus is shown for several Brassicaceae members. The position of the genomic region in the respective genome assembly is given underneath the species name in million base pairs (Mb). Grey curved beams connect the identified syntenic genes. The rectangle shaped arrows represent annotated genes. Genes located on the forward strand are shown in grey and genes located on the reverse strand are shown in black. *MYB24* homologs are marked in red and connected by light red lines. The assembly continuity at the *MYB24* locus was too low to analyse local synteny in *C. maritima, E. vesicaria*, and *C. hispanica*. A second *S. alba* locus sharing the same degree of local synteny is not shown for clarity (Additional file 6).

### Gene expression analyses of SG19 MYBs

Analysis of tissue-specific expression patterns of SG19 members in *B. napus* revealed that all *BnaMYB21* homologs are strongly expressed in stamens, pistils, sepals, and petals (Table 2). However, *BnaMYB21-2_A09p002640* is expressed at higher levels in roots and seed coat 21-28 DAF compared to the other *BnaMYB21* homologs. While the expression of *BnaMYB57* homologs, if expressed, in stamens and sepals was lower compared to *BnaMYB21* homologs, it was frequently higher in petals and pistils. Interestingly only *BnaMYB57-3_C03p034120* and *BnaMYB57-4_A03p028260* were expressed in all four floral tissues with *BnaMYB57-3* being exceptionally strongly expressed in petals. The *BnaMYB57-2_A05p000230* gene is expressed in pistils, sepals and petals but is only marginally expressed in stamens, while *BnaMYB57-1_C05p047780* is only expressed in petals. Interestingly, *BnaMYB57-4_A03p028260* revealed uniquely high expression in young seeds, while *BnaMYB57-3_C03p034120* showed uniquely high expression in seed coat 42 DAF and endosperm. To summarize, the expression patterns of *BnaMYB57-1_C05p047780* and *BnaMYB57-2_A05p000230* overlap completely with the other *BnaMYB57* homologs, which show as well similar expression patterns. Co-expression analysis of the majority of SG19 members in *B. napus* revealed a correlation level too low to be considered as strong co-expression. However, *BnaMYB57-3_C03p034120* and *BnaMYB57-4_A03p028260* were co-expressed with each other (Additional file 5).

**Table 2:** Tissue-specific expression of SG19 MYBs in *B. napus*. The tissue-specific expression of the identified *MYB21* and *MYB57* homologs in *B. napus* is presented in mean transcripts per million (TPMs). The number of analysed data sets per tissue is stated in brackets (n=X). Intensity of the blue colouration indicates the expression strength (darker = stronger expression). Abbreviations: weeks after pollination (WAP), days after pollination (DAP), days after flowering (DAF), days (D), shoot apical meristem (SAM).

|  | *MYB21-1*<br>*C09p003180* | *MYB21-2*<br>*A09p002640* | *MYB21-3*<br>*C07p017730* | *MYB21-4*<br>*A06p033190* | *MYB57-1*<br>*C05p047780* | *MYB57-2*<br>*A05p000230* | *MYB57-3*<br>*C03p034120* | *MYB57-4*<br>*A03p028260* |
| --- | --- | --- | --- | --- | --- | --- | --- | --- |
| SAM (n=16) | 0.4 | 0.9 | 0.2 | 0.0 | 0.0 | 0.0 | 0.0 | 0.0 |
| Anther prophase 1 (n=12) | 0.0 | 0.0 | 0.0 | 0.2 | 0.0 | 0.0 | 0.0 | 0.0 |
| Anther bolting (n=6) | 0.0 | 0.0 | 0.0 | 0.0 | 0.0 | 0.0 | 0.0 | 0.0 |
| Anther flowering (n=4) | 0.0 | 0.0 | 0.0 | 0.4 | 0.0 | 0.0 | 0.6 | 0.9 |
| Stamen (n=1) | 115.5 | 106.0 | 263.0 | 199.3 | 0.0 | 0.4 | 21.2 | 43.0 |
| Ovule (n=1) | 0.2 | 0.0 | 0.2 | 0.1 | 0.0 | 0.1 | 1.2 | 0.9 |
| Pistil (n=3) | 38.6 | 73.6 | 86.2 | 90.5 | 0.0 | 112.3 | 132.6 | 139.9 |
| Sepal (n=1) | 134.3 | 134.3 | 181.7 | 90.8 | 0.0 | 4.9 | 14.0 | 4.6 |
| Petal (n=2) | 201.9 | 339.1 | 358.9 | 496.3 | 113.5 | 592.6 | 1,521.1 | 360.5 |
| bud (n=33) | 0.7 | 2.6 | 1.7 | 1.9 | 0.0 | 0.0 | 0.5 | 0.6 |
| Silique 10-20DAF (n=13) | 0.2 | 1.0 | 0.0 | 0.1 | 0.0 | 0.1 | 0.6 | 0.2 |
| Silique 25DAF (n=6) | 0.3 | 0.5 | 0.2 | 0.2 | 0.0 | 0.1 | 0.2 | 0.1 |
| Silique 30DAF (n=6) | 0.0 | 0.0 | 0.1 | 0.1 | 0.0 | 0.1 | 0.2 | 0.1 |
| Silique 40DAF (n=2) | 0.0 | 0.1 | 0.1 | 0.0 | 0.0 | 0.0 | 0.0 | 0.0 |
| Seed 2WAP (n=1) | 0.0 | 0.0 | 0.0 | 0.0 | 0.0 | 0.0 | 1.4 | 5.6 |
| Seed 4WAP (n=1) | 0.0 | 0.0 | 0.0 | 0.1 | 0.0 | 0.1 | 0.3 | 0.3 |
| Seed 6WAP (n=1) | 0.0 | 0.0 | 0.0 | 0.0 | 0.0 | 0.0 | 0.1 | 0.1 |
| Seed 8WAP (n=1) | 0.1 | 0.0 | 0.0 | 0.0 | 0.0 | 0.0 | 0.0 | 0.0 |
| Seed brown 26DAF (n=1) | 1.2 | 1.1 | 0.2 | 0.3 | 0.0 | 0.1 | 0.3 | 0.1 |
| Seed yellow 26DAF (n=1) | 0.4 | 0.3 | 0.0 | 0.2 | 0.0 | 0.1 | 0.3 | 0.1 |
| Seed coat 14DAF (n=7) | 0.0 | 0.1 | 0.1 | 0.0 | 0.0 | 0.0 | 0.6 | 1.5 |
| Seed coat 21DAF (n=6) | 0.0 | 4.1 | 0.3 | 1.2 | 0.0 | 0.0 | 0.8 | 1.5 |
| Seed coat 28DAF (n=6) | 0.0 | 7.3 | 0.6 | 2.0 | 0.0 | 0.0 | 0.9 | 0.4 |
| Seed coat 35DAF (n=6) | 0.0 | 0.7 | 0.3 | 0.3 | 0.0 | 0.0 | 1.4 | 0.4 |
| Seed coat 42DAF (n=6) | 0.0 | 0.2 | 1.0 | 0.1 | 0.0 | 0.0 | 5.3 | 1.9 |
| Embryo (n=6) | 0.0 | 0.0 | 0.0 | 0.0 | 0.0 | 0.0 | 0.0 | 0.0 |
| Endosperm (n=8) | 0.0 | 0.0 | 1.0 | 0.5 | 0.0 | 0.0 | 3.5 | 1.1 |
| Seedling (n=9) | 0.0 | 0.6 | 0.0 | 0.0 | 0.0 | 0.1 | 0.5 | 0.1 |
| Cotyledon 7-10D (n=34) | 0.0 | 0.1 | 0.0 | 0.0 | 0.0 | 0.0 | 0.9 | 0.4 |
| Leaf juvenile (n=12) | 0.0 | 0.2 | 0.0 | 0.0 | 0.0 | 0.0 | 0.3 | 0.6 |
| Leaf old (n=12) | 0.0 | 0.1 | 0.1 | 0.0 | 0.0 | 0.0 | 1.1 | 1.2 |
| Internode flowering (n=6) | 0.0 | 0.1 | 0.1 | 0.0 | 0.0 | 0.1 | 0.0 | 0.1 |
| Stem (n=19) | 0.3 | 0.6 | 0.3 | 0.3 | 0.0 | 0.2 | 0.3 | 0.4 |
| Shoot (n=2) | 0.0 | 0.0 | 0.0 | 0.0 | 0.0 | 0.0 | 0.0 | 0.2 |
| Shoot apices (n=2) | 0.0 | 0.1 | 0.1 | 0.0 | 0.1 | 0.0 | 0.0 | 0.0 |
| Root seedling (n=13) | 0.0 | 0.0 | 0.0 | 0.0 | 0.0 | 0.0 | 0.0 | 0.2 |
| Root 30DAP (n=20) | 1.0 | 6.2 | 0.5 | 0.2 | 0.0 | 0.1 | 0.1 | 0.1 |
| Root 60DAP (n=2) | 0.6 | 0.4 | 0.1 | 0.0 | 0.0 | 0.0 | 0.0 | 0.0 |

## Discussion

In this study we analyzed flavonol regulators across 31 Brassicaceae species spanning 17 tribes. We identified a deep duplication giving rise to *MYB12, MYB111* and *MYB21* likely preceding the divergence of Brassiceae, which was followed by the loss of *MYB11* and *MYB24* after the divergence of the Brassiceae (Figure 8).

**Figure 8:**
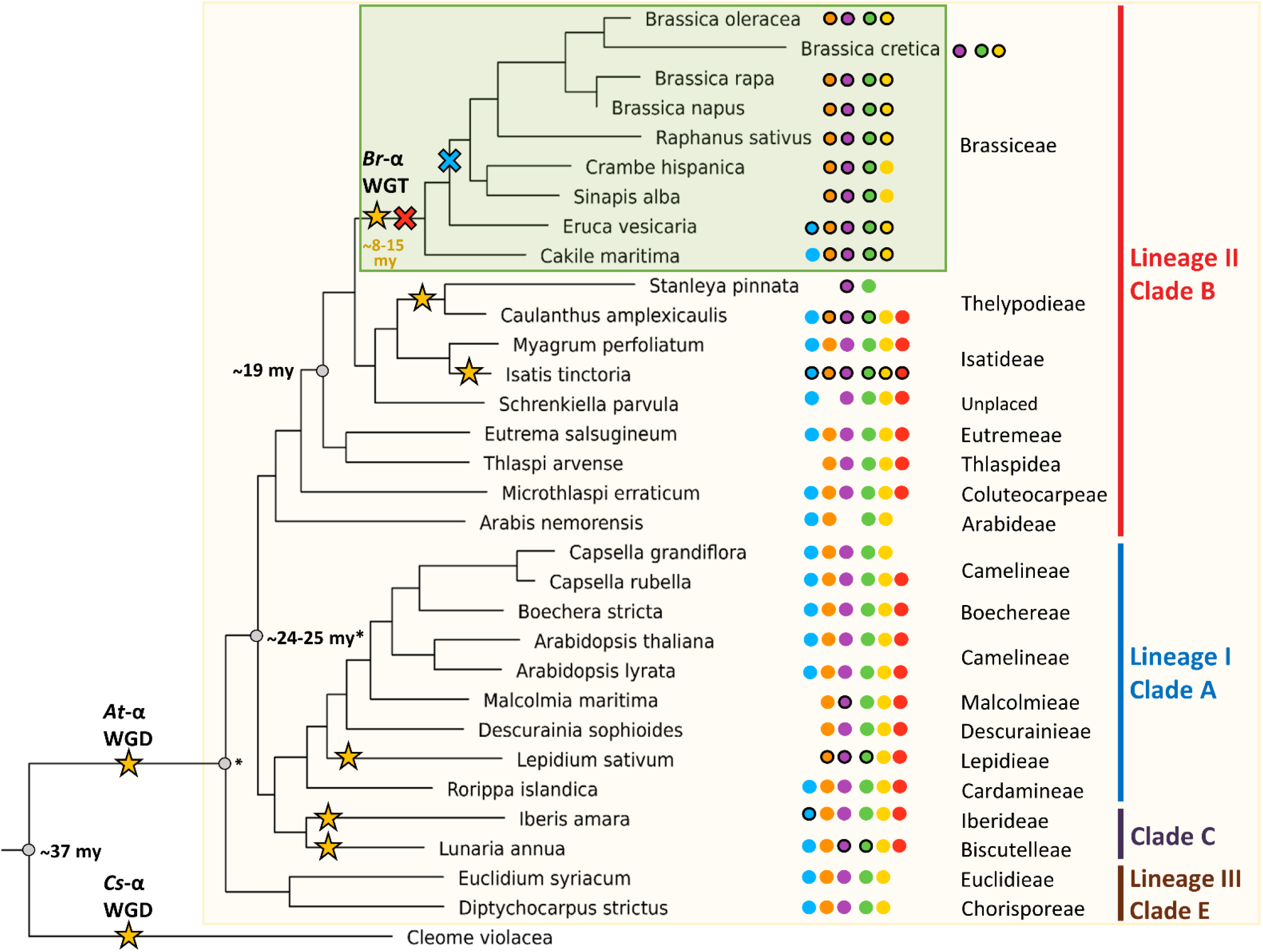
Graphical abstract of SG7 and SG19 evolution in Brassicaceae. The proposed duplication and gene loss events inside the Brassiceae are shown. SG7 and SG19 homologs identified in Brassicaceae species are marked with different coloured circles: *MYB11* in light blue, *MYB12* in orange, *MYB111* in violet, *MYB21* in green, *MYB57* in yellow, and *MYB24* in red. If at least two homologs were detected in the species the circle was marked with a dark outline. The assumed loss of *MYB11* is marked with a light blue cross, while the proposed loss of *MYB24* is marked with a red cross. The duplication events of *MYB12, MYB111* and *MYB21* likely preceded the divergence of the Brassiceae tribe.

### Polyploidization events have shaped the evolution of the SG7 and SG19 MYBs inside the Brassicaceae

WGD events are known to influence genetic diversification and species radiation. Polyploidization events allow an adaptive advantage by providing the genetic basis for gene neo- and subfunctionalisation [20]. Additionally, affected genomes are characterized by extensive re-diploidization, typically associated with chromosomal rearrangements, genome size reduction and increased fractionation [55]. These events can lead to gene losses while duplicated genomic regions can still be identified [56, 55]. Besides the paleo-polyploidization events At-ɣ, At-β, and At-α, lineage-specific meso-polyploidization events took place during the evolution of several Brassicaceae tribes including Brassiceae, Isatideae, and Thelypodieae [21, 22, 57, 23]. The meso-polyploidization event of *Isatis tinctoria* (Isatideae) likely resulted in the duplication of all SG7 and SG19 members as inferred by the close phylogenetic relationship of the duplicated homologs (Figure 4, Figure 6). These duplication events are thus independent from the observed duplication events inside the Brassiceae and Thelypodieae. The duplicated *MYB12, MYB111*, and *MYB21* homologs of the Thelypodieae fall into separate clades, thus suggesting that these duplication events might not be associated with the independent meso-polyploidization event but rather belong to a deeper duplication that took place in the common ancestor of Brassiceae and Thelypodieae. One of the most recent Brassicaceae phylogenies suggests Brassiceae and Thelypodieae to be closely related monophyletic sister clades while Isatideae is sister to both, supporting this hypothesis [4]. However, additional research including more data from Brassiceae sister tribes, e.g. the Sisymbrieae, is needed to further pin-point the timepoint of the *MYB12, MYB111*, and *MYB21* duplication events. The *MYB57* duplication observed in 7/9 Brassiceae species, but not in the Thelypodieae, is likely associated with the Brassiceae-specific whole-genome triplication (WGT) dated to 7.9–14.6 my [15, 16]. This Br-α WGT event was shown to have been followed by taxon- and lineage-specific chromosome rearrangements resulting in chromosome number reductions [15, 16], which might be associated with the observed secondary loss of one *MYB57* homolog in the closely related *Sinapis alba* and *Crambe hispanica* (Figure 6).

Succeeding these duplication events we identified the loss of *MYB11* after the divergence of *Eruca vesicaria* (Brassiceae) and the loss of *MYB24* after the divergence of the Brassiceae (Figure 4). The loss of *MYB11* and *MYB24* inside the Brassiceae was further supported by the absence of these homologs in the respective genomic regions showing the highest local synteny to the *MYB11* and *MYB24* loci in *A. thaliana* and other Brassicaceae species (Figure 5, Figure 7). Recently, Li *et al*. 2020 analysed the distribution of R2R3-MYBs in nine Brassicaceae (*A. thaliana, Arabidopsis lyrata, Capsella rubella, Capsella grandiflora, Boechera stricta, B. napus, B. oleracea, B. rapa, Eutrema salsugineum*) and seven non-Brassicaceae species (*Carica papaya, Theobroma cacao, Gossypium raimondii, Citrus clementina, Citrus sinensis, Manihot esculenta, Eucalyptus grandis*) [32]. In accordance with our results no *MYB11* or *MYB24* homolog was identified for the three analysed Brassiceae species and at least two *MYB12, MYB21, MYB111*, and *MYB57* homologs were detected for *B. rapa* and *B. napus*. However, for *B. oleracea* only one *MYB12, MYB111*, and *MYB21* homolog was identified, along with two *MYB57* homologs. This difference might be explained by the use of a short-read assembly (N50 = ∼27 kbp, 5,425 contigs) vs. a long-read assembly (N50 = ∼9,491 kbp, 264 contigs) used in this study in which more homologs could be resolved. In summary, the duplications of *MYB12, MYB111*, and *MYB21* identified in all Brassiceae species are derived from a deep duplication event presumably preceding the divergence of Brassiceae. The subsequent loss of *MYB24* and *MYB11* inside the Brassiceae might have occurred during the course of post-mesopolyploidization of the Br-α WGT event.

### The impact of gene redundancy and different tissue-expression pattern on SG7 and SG19 MYB evolution inside Brassicaceae

Gene redundancy accompanied with differential spatial expression has been observed for the SG7 MYBs in *A. thaliana* seedlings: *MYB12* is expressed in roots, while *MYB111* is expressed in cotyledons and *MYB11* is only marginally expressed in defined narrow domains of the seedling like the root tip and the apex of cotyledons [41]. Thus, *MYB12* and *MYB111* were designated as the main flavonol regulators in *A. thaliana* seedlings [41]. Moreover, Stracke *et al*. postulated that MYB12 and MYB11 regulate different targets involved in the production of specific flavonol derivatives because the single mutants displayed differences in the composition of flavonol derivatives. In contrast, the *MYB11* single mutant revealed a flavonol composition that is comparable to the wild type [41]. Moreover, the expression pattern of SG7 members in *B. napus* differs from the ones described for *A. thaliana* seedlings: *BnaMYB12* are predominantly expressed in reproductive tissues and *BnaMYB111* in anthers and buds. One of the main target genes of the SG7 members, *flavonol synthase* (*FLS*), is also mainly expressed in reproductive tissues in *B. napus* [58] indicating the relevance of the transcriptional activation of flavonol accumulation in reproductive tissues. Reduced flavonol levels were linked with decreased pollen viability and germination, as e.g. pollen germination increased with increasing flavonol concentrations and kaempferol supplementation rescued pollen fertility [59, 60]. In general, overlapping expression patterns of *BnaMYB12* and *BnaMYB111* homologs were identified, accompanied by tissue-specific expression of single *BnaMYB12* and *BnaMYB111* homologs. The majority of *BnaMYB12* and *BnaMYB111* homologs were co-expressed with genes involved in or associated with flavonoid biosynthesis, indicating their proposed role in the regulation of this pathway. These findings indicate that the *BnaMYB12* and *BnaMYB111* homologs might be active in the same tissues showing (partial) functional redundancy, while the unique expression domains of single homologs could explain why single homologs are retained. Additionally, specific sequence features might play a role in subfamily and gene retention, as *Bna*R2R3-MYB subfamilies with a specific intron pattern are more likely to be retained [30, 32]. The *BnaMYB21* and *BnaMYB57* homologs revealed strong and overlapping expression in stamens, pistils, sepals and petals. Again tissue-specific expression of single *BnaMYB21* and *BnaMYB57* homologs was identified. Taken together, it seems likely that the duplicated *MYB12* and *MYB111* homologs and *MYB21* and *MYB57* homologs inside the Brassiceae can compensate for the loss of *MYB11* and *MYB24*, respectively. Recent functional analyses of *BnaWER* homologs (S15) indicate that genes derived from the same subfamily, which share high sequence similarity and similar expression patterns, frequently show functional redundancy [32]. However, further research is necessary to elucidate the biological meaning and function of the *MYB12, MYB111, MYB21*, and *MYB57* duplications and proteins, respectively.

One well-known example of the evolution of novel traits in the Brassicales, including Brassicaceae, is the emergence of glucosinolates (GSLs) along with the corresponding R2R3-MYB transcriptional regulators *MYB28, MYB29, MYB34, MYB51, MYB76* and *MYB122*, which belong to subgroup 12 [25, 61]. This MYB clade is proposed to result from the At-β paleo-polyploidization event [62]. *MYB28, MYB29*, and *MYB76* act as positive regulators of aliphatic GLSs with overlapping functions and *MYB28* and *MYB29* as main regulators being able to compensate the lack of *MYB76* in *A. thaliana* [63]. While *MYB76* is present in *A. thaliana* (Camelineae), no *MYB76* has been identified in *Brassica* species (Brassiceae) [61] posing a striking example of gene loss inside specific Brassicaceae species. Interestingly, we observed that the divergence of *MYB11* and *MYB12*, as well as *MYB21* and *MYB24*, likely occurred after the divergence of the Cleomaceae from its sister group the Brassicaceae (Figure 3). Previous studies included only *A. thaliana* as a single Brassicaceae species [30, 31], thus could not analyse Brassicaceae-specific expansion of SG7 and SG19 MYBs. However, Li *et al*. 2020 investigated the SG7 and SG19 homologs of nine Brassicaceae species and seven non-Brassicaceae species, thereby revealing five Brassicaceae-specific subfamilies and five subfamilies which were absent from the investigated Brassicaceae species [32]. In accordance with our hypothesis, the non-Brassicaceae SG7 and SG19 homologs did not fall into two separate *MYB11* and *MYB12* clades, as well as *MYB21* and *MYB24* clades, respectively, while the Brassicaceae homologs did [32]. Thus our study used a broad range of Brassicaceaeand related species like *Cleome violacea*, allowing the in-depth analysis and identification of Brassicaceae-specific expansion of SG7 and SG19 MYBs. This finding serves as an example of the adaptive evolution of the flavonol-regulating R2R3-MYB transcription factors frequently accompanied by sub- and neofunctionalization in Brassicaceae species where a *MYB11* and *MYB24* homolog was retained. Moreover, our results suggest that lineage-specific expansion or reduction of MYB subfamilies might have occurred frequently in the Brassicaceae, in line with the high degree of flexibility and complex evolution observed for the *B. napus* R2R3-MYB subfamilies.

### Limitations of the study

The quality of the sequence data sets used in this study varies between species. Different degrees of completeness can influence the identification of homologs. For example, no *MYB11, MYB12, MYB24*, and *MYB57* homolog was identified in *Stanleya pinnata*, probably due to the low completeness (71 % complete BUSCOs) observed for this data set (Additional file 4). Additionally, *Brassica cretica* revealed a comparably low completeness of 74.5 % and no *MYB12* homolog was identified (Additional file 4). The recent release of genomic resources for several Brassicaceae members allowed us to investigate the evolution of the SG7 and SG19 MYBs in great detail. Thus, in this study we were able to cover 17 of the 51 Brassicaceae tribes with at least one representative species. However, additional genome sequences of Brassicaceae species will help to support our hypotheses and to further narrow down the time-point of the SG7 and SG19 duplication and gene loss events. The species tree revealed minor differences to the phylogeny of taxonomic studies like Huang et al. 2015 [9], Nikolov *et al*., 2019 [10] and Walden *et al*. 2020 [4]. However, the phylogenetic positions of the tribes is still not fully resolved due to different results derived from nuclear and plastid data which, among other reasons, explains the inconsistencies of Brassicaceae taxonomy studies (summarised in Walden *et al*., 2020).

## Conclusions

In this study we unraveled the evolution of the flavonol regulators SG7 and SG19 R2R3-MYBs in the Brassicaceae with focus on the tribe Brassiceae (Figure 8). A deep duplication of the SG7 MYBs *MYB12* and *MYB111*, likely preceding the divergence of Brassiceae, was followed by the loss of *MYB11* after the divergence of *E. vesicaria*. Similarly, a duplication of *MYB21* likely preceding the divergence of the Brassiceae was associated with the loss of *MYB24* inside the Brassiceae. Due to the overlapping spatio-temporal expression patterns of the SG7 and SG19 members in the Brassiceae member *B. napus*, the loss of *MYB11* and *MYB24* is likely to be compensated for by the remaining homologs. Therefore, we propose that polyploidization events and gene redundancy have influenced the evolution of the flavonol regulators in the Brassicaceae, especially in the tribe Brassiceae.

## Methods

### Data collection, quality control and species tree generation

Genomic data sets of 44 species, including 31 species of the Brassicaceae, were retrieved mainly from Phytozome, NCBI and Genoscope (Additional file 1). To assess the completeness and duplication level of all annotated polypeptide sequences BUSCO v3.0.2 was deployed using the embryophyta_odb9 lineage dataset in protein mode [64]. OrthoFinder v2.5.4 [65–67] was used to construct a species tree using the 44 proteome data sets as input.

### Genome-wide identification of MYB homologs

Genome-wide identification of MYB and MYB-like transcription factors was performed using MYB annotator v0.153 [68]. MYB annotator was run with the default bait sequences and the proteome data sets of all 44 species were subjected to this analysis. The extracted MYB polypeptide sequences per species were combined and used for the phylogenetic analysis.

### Phylogenetic tree construction

For the generation of a phylogenetic tree, first the full-length polypeptide sequences of the genome-wide identified MYB homologs per species were combined into one file (Additional file 7) and then used for the construction of a MAFFT v7.475 [69] alignment. This analysis covered 44 species (Additional file 1). Next, a codon alignment was produced via pxaa2cdn [70] i.e. converting the amino acids of the alignment back to their respective codons. As no CDS file was available for *Arabis nemorensis, Brassica cretica* and *Microthalspi erraticum*, these species were not incorporated in this analysis. However, the SG7 and SG19 homologs identified in these species based on polypeptide sequences are listed in Additional file 8. Subsequently, the alignment was cleaned by removal of all columns with less than 10 percent occupancy as described before [71]. The cleaned alignment was then used for the construction of an approximately-maximum-likelihood phylogenetic tree constructed with FastTree 2 [72] using the WAG model and 10,000 bootstrap replications in addition to the following parameters to increase accuracy: -spr 4 -mlacc 2 -slownni -gamma. This phylogenetic tree covering all genome-wide MYBs from 41 species was then used for the identification of the SG7 and SG19 clade followed by the extraction of the included MYB polypeptide sequences by a customized python script (extract_red.py) [73]. Additionally, the SG5 and MYB99 homologs were extracted because MYB123 (SG5) regulates a competing branch of the flavonoid pathway and is sister clade to SG7 and MYB99 is involved in the regulation of SG19 MYBs. Again, an alignment of polypeptide sequences (corresponding CDS sequences are listed in Additional file 9) was constructed followed by its conversion into a codon alignment and cleaning as described above. Next, the cleaned codon alignment was used to construct a tree via RAxML-NG v.1.0.1 [74] using the GTR+GAMMA model. The best-scoring topology was inferred from 50 tree searches using 25 random and 25 parsimony-based starting trees. To infer a bootstrap tree, again the GTR+GAMMA model was used including 9800 bootstrap replicates until bootstrap convergence was reached after 8750 bootstraps (weighted Robinson-Foulds (RF) distance = 0.646, 1% cutoff). The bootstrap support values were then mapped onto the best-scoring Maximum Likelihood (ML) tree. After monophyletic tip masking, the resulting tree with bootstrap support values was visualized using FigTree v1.4.3 (Additional file 3). MYBs per species were classified according to their relationships with *A. thaliana* homologs.

### Synteny and BLAST analysis

JCVI [75] was used to analyse local synteny and visualize syntenic regions. To analyse a potential gene loss event in a species in detail a TBLASTN [76] against the high local synteny regions using *Ath*MYB11 and *Ath*MYB24 as queries was performed with all Brassiceae members, *I. tinctoria* and *M. perfoliatum*. Moreover, TBLASTN was run against the respective assemblies of these species to search for potential gene fragments of *MYB11* and *MYB24* outside of the syntenic regions. For this analysis a customized python script was used (TBLASTN_check.py) [73], which identifies whether a TBLASTN hit is located inside an annotated gene or not. If several blast hits correspond to the same gene (e.g. multiple exons), the identifier of this gene will only be extracted once. If the TBLASTN hit is not located inside a gene, the start and end position on the subject sequence will be extracted and used for a web-based BLASTN search to identify potential homologs. The top five hits were then used to extract the amino acid sequence from the corresponding gene ID and then subjected to phylogenetic analysis including all 126 *Ath*R2R3-MYBs via FastTree 2 [72]. This analysis revealed their closest *Ath*MYB homolog for classification. If the closest homolog was not MYB11 or MYB24, this would further support the absence of these homologs in the analysed species.

### Gene expression analysis

Public RNA-Seq data sets were used and retrieved from the Sequence Read Archive via fastq-dump v.2.9.64 [77] to analyze the expression of MYB genes across various tissues (Additional file 10). Transcript abundance, i.e. read counts and transcripts per millions (TPMs), was calculated via kallisto v. 0.44 [78] using default parameters and the transcript file of the *B. napus* cultivar Express 617 [79]. The heatmap was constructed with a customized python script calculating mean TPMs per tissue using 276 paired-end RNA-Seq data sets from *B. napus* as previously described [58]. Condition-independent co-expression analysis was performed as described before [58] to identify co-expressed genes using Spearman’s correlation coefficient by incorporating 696 *B. napus* RNA-Seq data sets.

## Supplementary Information

**Additional file 1: Information about the used data sets**. The version and reference of the data set per species are listed. Moreover, the completeness and duplication level of the respective proteome data set per species is stated based on BUSCOs. Brassicaceae species are highlighted in green.

**Additional file 2: 1R-, R2R3-, and 3R-MYBs composition per analysed species**. The number of 1R-, R2R3-, and 3R-MYBs per analysed species is listed as identified and classified by MYB annotator. Brassicaceae species are highlighted in green. Brassiceae species are shown in italics.

**Additional file 3: Phylogenetic tree of SG5, SG7, SG19 and MYB99 members**. Bootstrap values are represented as percentages.

**Additional file 4: Number and gene identifiers of the identified SG7 and SG19 homologs per Brassicaceae species**. Brassiceae species are highlighted in green. If more than one SG7 and SG19 homolog was identified the number was marked in bold. The BUSCO completeness of the data set per species is stated.

**Additional file 5: Co-expression analysis of SG7 and SG19 MYB family members in *B. napus***. Yellow highlighted homologs are described in the main text and the threshold for strong co-expression (Spearman’s correlation coefficient >= 0.7) is marked with a black line.

**Additional file 6: Synteny analysis of the *MYB24* locus including the second *S. alba* high local synteny locus. Additional file 7: Polypeptide sequences from the genome-wide MYBs identified by MYB annotator**.

**Additional file 8: Polypeptide sequences of SG7 and SG19 homologs identified in *Arabis nemorensis, Brassica cretica*, and *Microthalspi erraticum* via MYB annotator**.

**Additional file 9: CDS sequences from the phylogenetic tree of SG5, SG7, SG19 and MYB99 members**.

**Additional file 10: SRA data sets used for tissue-specific RNA-Seq analysis**. The number of analysed data sets per tissue is stated in brackets (n=X). The heatmap from white via light to dark blue indicates the expression strength with dark blue symbolizing high expression. Abbreviations: weeks after pollination (WAP), days after pollination (DAP), days after flowering (DAF), days (D), shoot apical meristem (SAM).

## Declarations

## Acknowledgments

We are grateful to all researchers who submitted the underlying sequences to the appropriate databases, and published their experimental findings. Some of the sequence data sets used were produced by the US Department of Energy Joint Genome Institute. We thank the Center for Biotechnology (CeBiTec) at Bielefeld University for providing an environment to perform the computational analyses.

## Funding

We acknowledge support for the publication costs by the Open Access Publication Fund of Bielefeld University and the Deutsche Forschungsgemeinschaft (DFG), as well as the support of the German Academic Exchange Service.

## Availability of data and materials

All datasets underlying this study are publicly available or included within the additional files.

## Authors’ contributions

HMS and BJG designed the research. HMS performed bioinformatic analyses. HMS and BJG interpreted the results and wrote the manuscript. Both authors read and approved the final version of the manuscript.

## Ethics approval and consent to participate

Not applicable.

## Consent for publication

Not applicable.

## Competing interests

The authors declare that they have no competing interests.

## Notes

### Competing Interest Statement

The authors have declared no competing interest.

