## Supplementary figures and images for "Analysis of flavonol regulator evolution in the Brassicaceae reveals *MYB12, MYB111* and *MYB21* duplications associated with *MYB11* and *MYB24* gene loss"

### Additional file 3

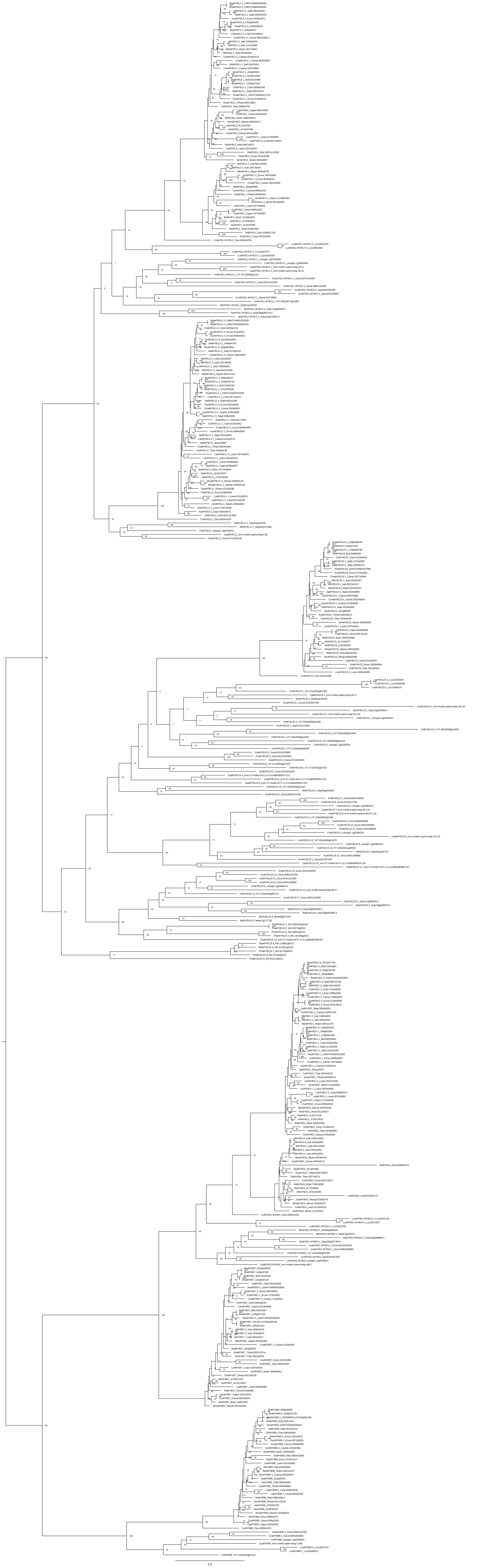

### Additional file 6

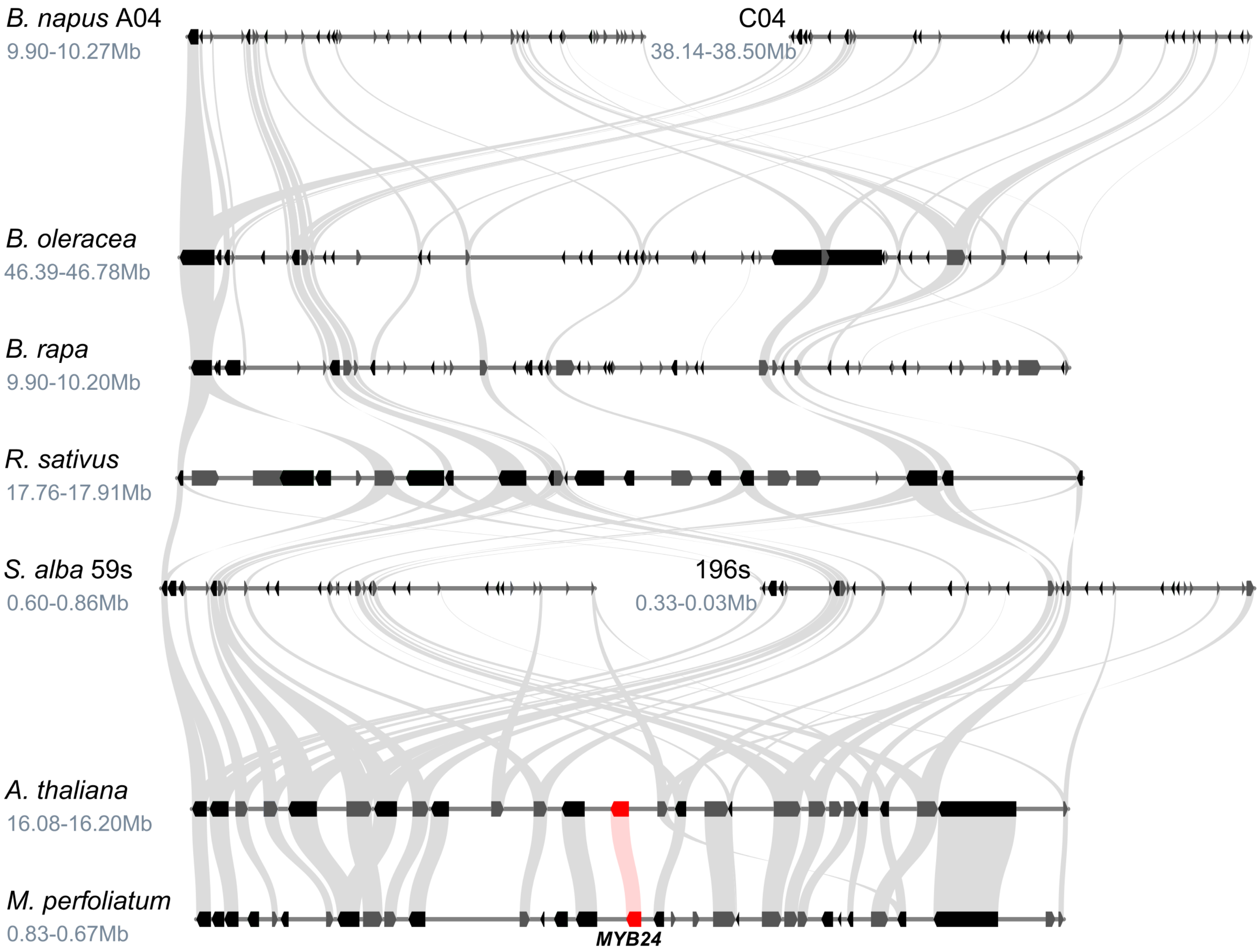
